# Angle-Tuned TMS Coils: Building Blocks for Brain Stimulation with Improved Depth-Spread Performance

**DOI:** 10.1101/2021.12.13.472217

**Authors:** Hedyeh Bagherzadeh, Qinglei Meng, Zhi-De Deng, Hanbing Lu, Elliott Hong, Yihong Yang, Fow-Sen Choa

**Affiliations:** Department of Computer Science and Electrical Engineering, University of Maryland, Baltimore County, MD, USA; Neuroimaging Research Branch, National Institute on Drug Abuse, Intramural Research Programs, National Institutes of Health, Baltimore, MD, USA; Noninvasive Neuromodulation Unit, National Institute of Mental Health, National Institutes of Health, Bethesda, MD, USA; Maryland Psychiatric Research Center, Department of Psychiatry, University of Maryland School of Medicine, Baltimore, MD, USA

**Keywords:** TMS, Electric Field, Stimulation Depth, Spread, Decay Rate

## Abstract

Coordinated whole-brain neural dynamics are essential for proper control of the functionality of different brain systems. Multisite simultaneous or sequential stimulations may provide tools for mechanistic studies of brain functions and the treatment of neuropsychiatric disorders. Conventional circular and figure-8 Transcranial Magnetic Stimulation (TMS) coils occupy a large footprint, and it is difficult to reach desired multiple stimulation locations with close proximity for comprehensive multisite stimulations. These conventional coils, limited by the depth-spread tradeoff rule, also lack the required focality for targeted stimulations. In this work, we propose and demonstrate angle-tuned TMS (AT) coils with an intrinsically reduced footprint with their geometric arrangements of stacking and angle tuning. The stimulation depth can be adjusted with the coil stacking number, and the field spread can be reduced by increasing the tilted wire-wrapping angle of the coils. With either smaller or larger diameter coils than a standard commercial figure-8 coil, we show, theoretically and experimentally, improved field decay rate and field intensity, and the reduced field spread spot size at different stimulation depths. These results indicate that the proposed novel coil establishes a better depth-spread tradeoff curve than the conventional circular and figure-8 coils. This coil design has a simple and single element structure and provides a promising solution for an improved multisite brain stimulation performance and serves as the building block of more complex coils for further depth-spread improvements.

## 1. INTRODUCTION

Simple behavioral activities involve multiple brain networks, and each brain network covers multiple brain regions^1,2^. Therefore, multisite neuromodulation with controlled timing may provide a critical tool for mechanistic studies of brain functions and treatment of neurologic and psychiatric disorders^3–10^. One such neuromodulatory technique is transcranial magnetic stimulation (TMS), which applies electric field pulses to the brain via a stimulation coil placed over the scalp. Conventional circular and figure-8 TMS coils occupy a substantial footprint, defined here as the tangential surface area the coil occupies in the contact surface plane closest to the head. It is also difficult to reach the desired multiple stimulation locations with close proximity for comprehensive multisite stimulation with these types of coils. The prospect of multisite stimulation for rodents is even worse due to the smaller brain size and the difficulty to proportionally shrink the coil size and keep up with the required stimulation field strengths^11^.

Furthermore, at the frequency that TMS operates, the electric field invariably diverges and decays along the path as it reaches deeper brain regions^12^. There have been consistent efforts to reduce field divergence and increase field strength to reach deep brain regions with a small spread. Since field emission from TMS coils is low frequency and near-field, the coils’ geometric arrangement is crucial in shaping the emitted field distribution. A variety of TMS coil geometries have been adopted along with additional methods for field divergence improvement^13–28^. To optimize the design of TMS coil’s electric field distribution, theoretical analysis based on analytical models^12,29–32^ or numerical simulations using either finite element method (FEM) or finite difference method (FDM)^33–42^ have been implemented along with additional physical and experimental verifications^43–45^. In general, the coil performance is governed by a depth–spread tradeoff^27^. Larger aperture coils have a smaller field divergence and can reach deeper regions with a slower field decay rate; however, the field spread is already significant when initiated from the aperture. Smaller aperture coils have a more substantial field divergence; it quickly spreads with a faster field decay rate even though the field spread is initially small.

Geometric shaping of the field can also be accomplished by using multiple coils to achieve a less spread field distribution. As a result, the field distribution in the head has a more elliptical shape than just spherical, and deeper stimulation depth can be accomplished with a smaller tangential field spread. Based on such a concept, using coils with different sizes and polarities to trim the combined near field elements have been proposed^22,23,46,47^. Notably, in Meng et al. ‘s work, field-shaping using an array of passive rings has the advantage of simplicity, requiring only one power supply and less energy consumption^11^. However, the trimming is less controllable compared with shaping through active elements. Nevertheless, using active elements for field-shaping needs accurate control of counter field generations and cancellations with large and small coils to provide sufficient spatial spherical harmonic components; with this counter field design, more power consumption is required.

The importance of multisite stimulation of different brain regions has culminated in various coil designs for this purpose. De Lara et al.^48^ proposed a three-axis coil design for multichannel stimulation providing accurate electric field steerability and targeting, which can be used in concurrent TMS–fMRI studies. However, simultaneous control of multiple power supplies, each with a different setting, and the variation of mutual inductance among the coils can make the emitted field different from the simulations. Conducting measurements to calibrate the system for each application makes the procedures challenging to execute. Additionally, the stimulation depth is restricted by the smaller size coils. Koponen et al.^49^ designed a 5-coil multi-locus apparatus to accomplish controlling the stimulation location similar to the goals of De Lara et al.^48^. However, the designed composed of stacked layers of orthogonally-oriented figure-8, cloverleaf, and circular coils with a substantial tangential footprint. The steerable proportion of the induced field lies at the center of the stack and could not cover a larger cortex area. It also encountered similar power and calibration issues as those of De Lara et al.^48^.

In this work, novel angle-tuned (AT) ring coils are proposed to reduce the individual coil footprint and improve the depth–spread characteristics. The field-shaping technique and the structure are simple and do not require counter field generations. So, it is easy to implement and modify. Stacking multiple coils helps to enhance field strength and reduce the footprint. It also helps to increase the field penetration depth through the modification of the field’s geometric distribution. Increasing the coil wire-wrapping angle reduces the field spread at the price of sacrificing a little depth performance due to part of the coil being pulled away from the head surface. Below we present theoretical and experimental comparisons of a commercial figure-8 coil’s field emission distributions with multi-stacked and angle-tuned coils. These novel coils demonstrated better spread, higher induced electric field strength, better field decay rate, and smaller footprints than conventional coils.

## 2. METHOD & MATERIAL

The simulations of the coils with a human head model were implemented using COMSOL (COMSOL Multiphysics, Version 5.5). The head model^27^ is a homogeneous sphere with a 17 cm diameter and isotropic electrical conductivity of 0.33 S/m^-1^. Figure 1(a) illustrates our TMS coil design. The coil has a fixed winding width (the difference between the outer and inner diameters) of 1 cm. In the simulations, we varied the windings’ inner and outer diameters ranging from 1–99 cm and 2–100 cm, respectively. We also changed the coil stacking number from 2 to 5 and then to 9 along the central axis (Z-axis) with a tilting angle of up to 70 degrees with a total height of 12.0 cm, 21.8 cm, and 34.8 cm, respectively. The layers are connected in series; the current excitation in all coils is a sinusoidal wave with a frequency of 5 kHz. The coils are placed 0.5 cm away from the head surface.

**Figure 1:**
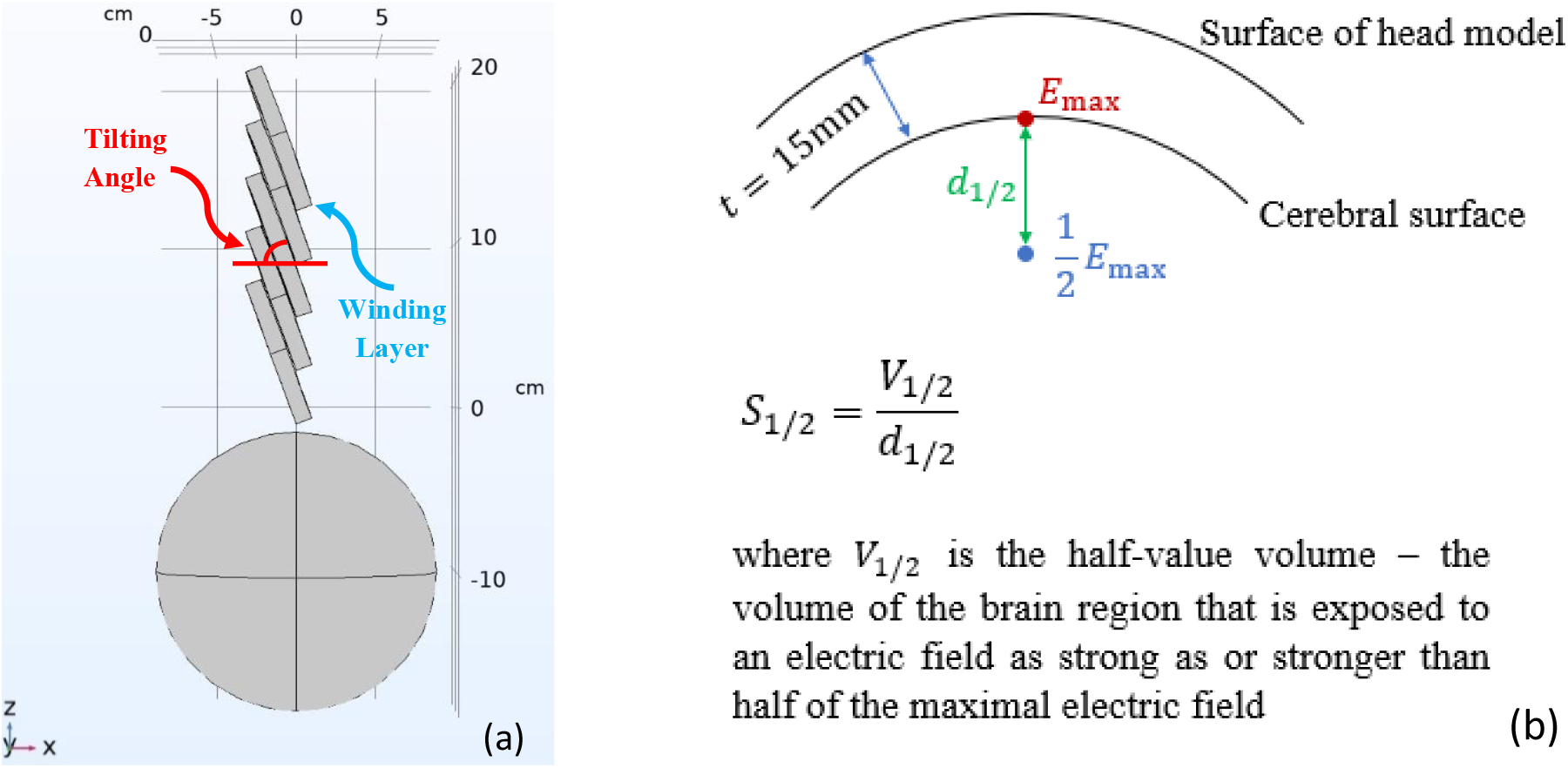
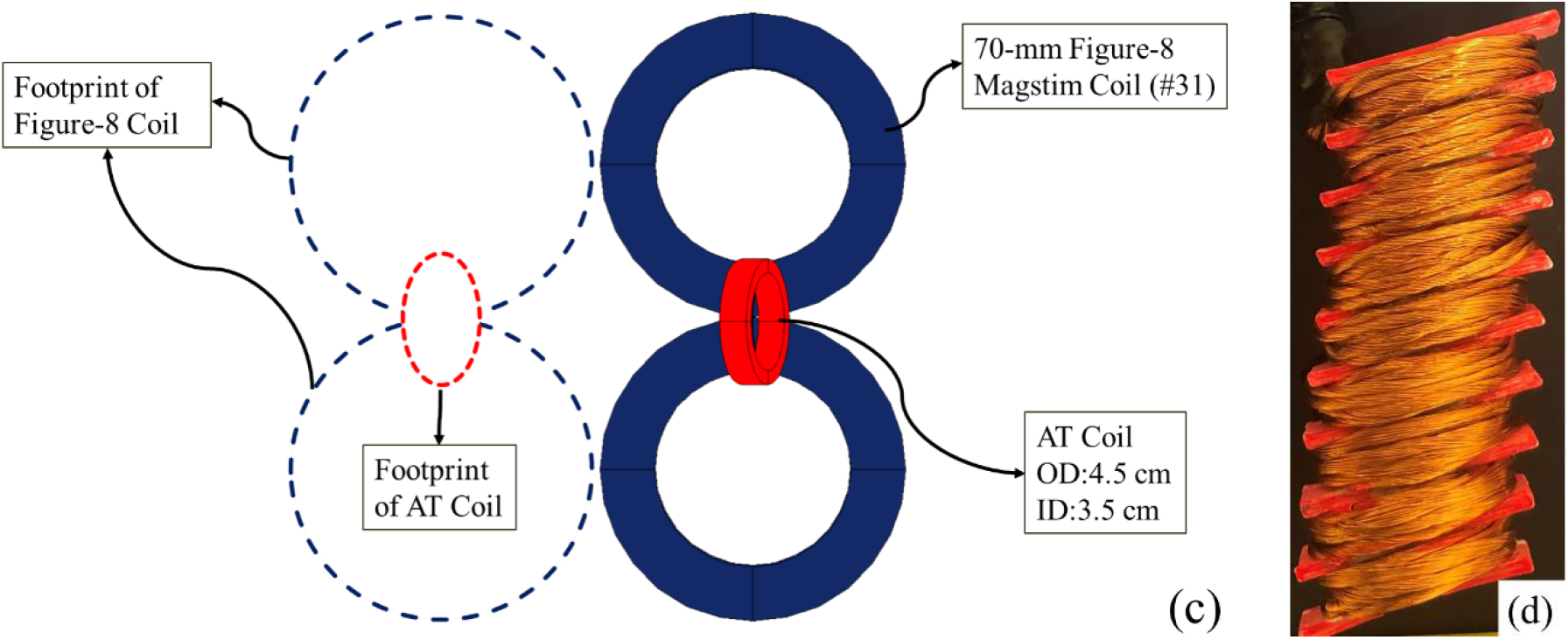
**a) Illustration of side-view for the proposed angle-tuned TMS coil** with five winding layers located above a homogenous and spherical head model, **b) The definitions of the half-value depth (d**_**1/2**_**), half-value volume (V**_**1/2**_**), and half-value spread (S**_**1/2**_) used for the estimation of the depth-spread characteristics for the proposed coils; The cerebral cortex is defined 1.5 cm below the surface of the head model. The half-value depth is defined as the radial distance from the cortical surface to the deepest point where the electric field strength is half of the maximum field strength on the cortical surface, **c) A plan view of the definition of the footprint**. The footprint is identified as the tangential surface area the coil occupies in the contact surface plane closest to the head, dictating the number of coils that can be simultaneously operated on the subject’s head. For the figure-8 coil (shown in blue), the footprint involves two circular coils with an outer diameter of 8.7 cm for each, resulting in a total footprint area of about 120 cm^2^ (dotted blue lines). In contrast, for the AT coil (shown in red), this value is calculated by multiplying the coil’s surface area by the cosine of the tilting angle. The footprint for a single 4.5-cm outer diameter is about 5.5 cm^2^ (dotted red lines), d) **Fabricated angle-tuned coil** for experimental measurements with nine winding layers wrapped over a 3-D printed coil holder. The fabricated coil has an inner and outer diameter of 1 cm and 3 cm, respectively, with a tilting angle of 20 degrees.

Figure 1(b) illustrates the definition of stimulation depth and spread in the head model introduced by the half-value depth (d_1/2_), half-value spread (S_1/2_), and half-value volume (V_1/2_) described in Deng et al.^27^. To compare our COMSOL simulations to previous studies, we selected three coils and used the same coil parameters as used in [27]. The half-value depth and spread of the three coils were analyzed: 50 mm and 70 mm circular coil (#1 and #4) and the 70 mm figure-8 (#31) Magstim coil. For all three cases, our simulated depth–spread results within 1% of previously reported^27^ results, as shown in Figure 2. We have also included other simulation results from Deng et al.^27^ in the background of the depth-spread plot as a reference forming the two best-fit curves for circular coils (solid line) and figure-8 coils (dashed line). We use the two curves as references to illustrate how different coil design parameters, such as tilting angle, number of winding layers, coil location, rotation angle, and outer diameter size, can affect the performance in terms of locations in the S_1/2_ vs. d_1/2_ plot. We further compare both simulation and experimental results of a commercial figure-8 coil and two of our angle-tuned coils with different coil diameters and tilting angles and show their field intensities and spot sizes at different distances from the coil.

**Figure 2:**
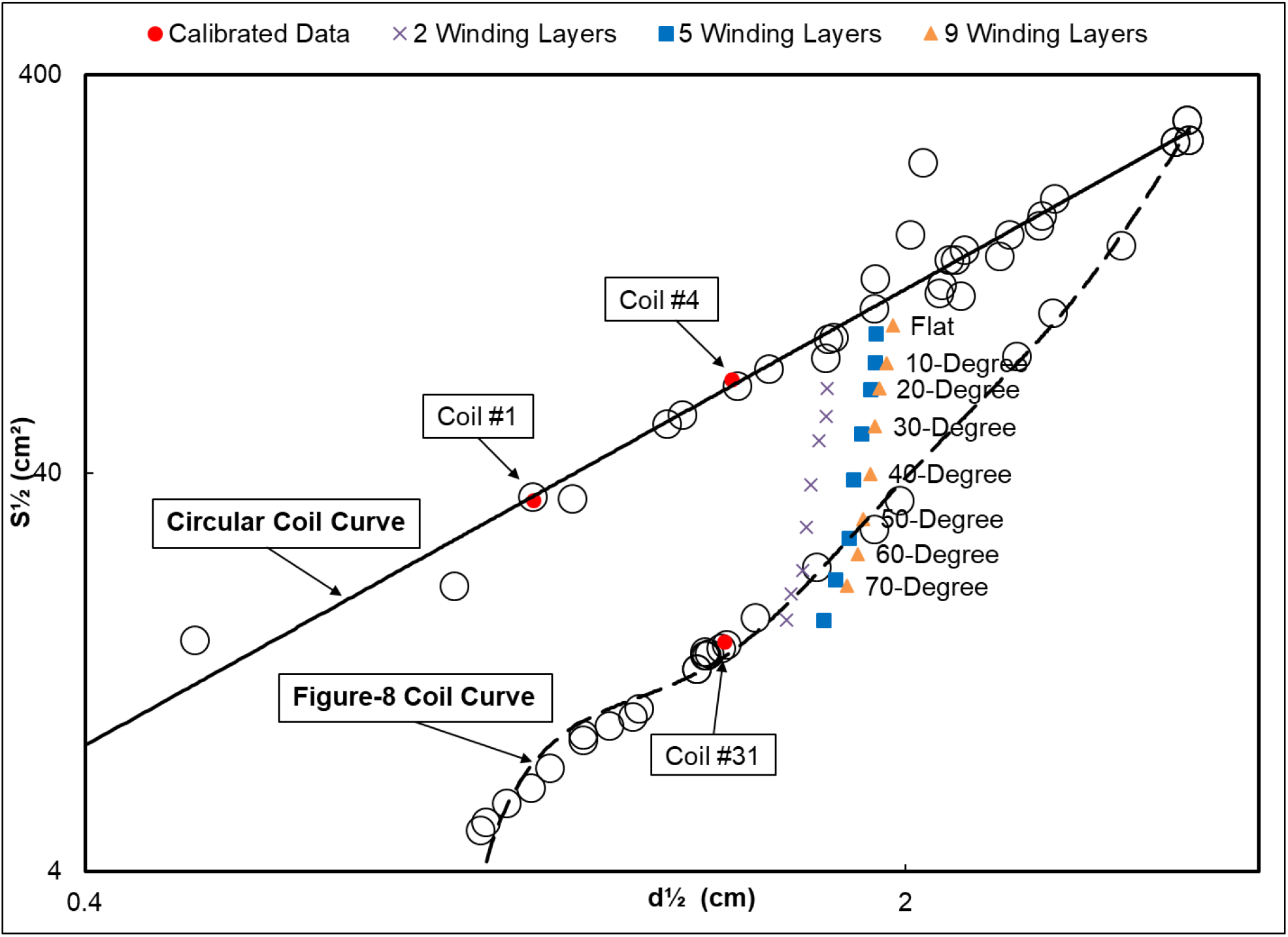
Effect of tilting angle and accumulation of wire wrapped coils on S_1/2_ and d_1/2_. The circles represent the locations of the previously studied coils by [27] with the solid and dashed lines showing the best-fit curves for the circular and figure-8 type of coils, respectively. The red dots are the calibrations performed in this study in COMSOL to validate the simulation technique. For all three cases, the calibration results are within 1% of the previously reported data. The AT coils have an inner and outer diameter of 8 cm and 9 cm, respectively, with tilting angles ranging from 0 to 70 degrees with 10-degree steps and winding layers of 2, 5, and 9.

Figure 1(c) demonstrates the definition of footprint in this work, characterized as the tangential surface area that the coil occupies in the projected surface plane. For example, the 70-mm figure-8 Magstim coil has a high lateral surface area of 120 cm^2,^ making it difficult to operate two coils on the human head at the same time. For the proposed AT coils, the footprint is calculated by multiplying the coil’s area by the cosine of the tilting angle. For the AT coil with an outer diameter of 4.5 cm, the footprint is about 5.5 cm^2^, approximately 95% less than the figure-8 coil (red dotted line for AT coil versus blue dotted line for figure-8 coil).

In our experimental works, angle tuned coils were fabricated using wire-wrapping over 3-D printed coil holders, which were printed with different angles and dimensions. The wires are made of Litz wire bundles with 135 pieces of insulated AWG30 wires for flexible bending and high current operations. The total number of turns for each coil was adjusted to be compatible with the inductance of the figure-8 coil (around 20 µH). Figure 1(d) shows the manufactured coil for the experimental measurements. A Magstim 200 (Magstim Co Ltd, Whitland, UK.) stimulator was used to drive these coils with the power set at 30% during the measurements. The coil’s electric field distribution was measured in a 5 mm step using calibrated high-spatial-resolution vector-field probes^50^. The total field strength, |E| = (E_x_^2^+E_y_^2^+E_z_^2^)^1/2^, is obtained from measuring the x, y, and z-direction field components at each step within the X-Y planes. We normalized the obtained values for easy comparison between experimental and simulation results.

## 3. RESULTS

### 3.1 FEM SIMULATIONS

Figure 2 illustrates the performance of the AT coils with an inner diameter of 8 cm and an outer diameter of 9 cm with tilting angles ranging from 0 to 70 degrees with 10-degree steps and stacking numbers of 2 (purple cross), 5 (blue square) and 9 (orange triangle). When the tilting angle is increased from 0 degrees (flat circular coil stack) to 70 degrees, the spread is significantly reduced with a small reduction in the depth. The reduction in the depth may be due to the tilting effect since part of the coil is pulling away from the head model.

The coils’ depth–spread performance surpasses the figure-8 coil curve when the tilting angle reaches about 50–60 degrees and beyond. This improvement clearly shows that the primary effect of the angle-tuning is to reduce the field spread. On the other hand, when the coil stacking number increases, the depth performance improves. This effect seems to be saturated from 5 to 9 coils. To check one numerical example of improvement, we compare the depth–spread of a five-winding-layer, 70-degree tilted coil with the 70-mm figure-8 Magstim coil (#31). They have about a similar spread, but the AT coil has a 20% deeper half-depth and has a smaller footprint by about 82%, as the figure-8 coil has a footprint of about 120 cm^2^ and the AT coil has a footprint of 22 cm^2^.

The proposed coils have a winding width of 1 cm, which is different from the coil width used in [27]. Reducing the winding width can slightly reduce the spread. So, S_1/2_-d_1/2_ location of the flat (zero-degrees tilting) AT coils in Figure 2 do not necessarily merge into the circular-coil tradeoff curve of [27]. Furthermore, since AT coils are tilted, we allow the coil and head model alignment not to be symmetric. The depth–spread improvement can also be the result of the position alignment. As shown in Figure 3(a), the position-1 alignment is more like a symmetric coil, where the central axis of the coil aligns with the Z-axis of the spherical head model. As shown in Figure 3(d), when the alignment changes, the depth–spread performance also changes. The position-2 arrangement, in which the coil’s deepest point is aligned with the Z-axis, provides the best depth– spread performance.

**Figure 3:**
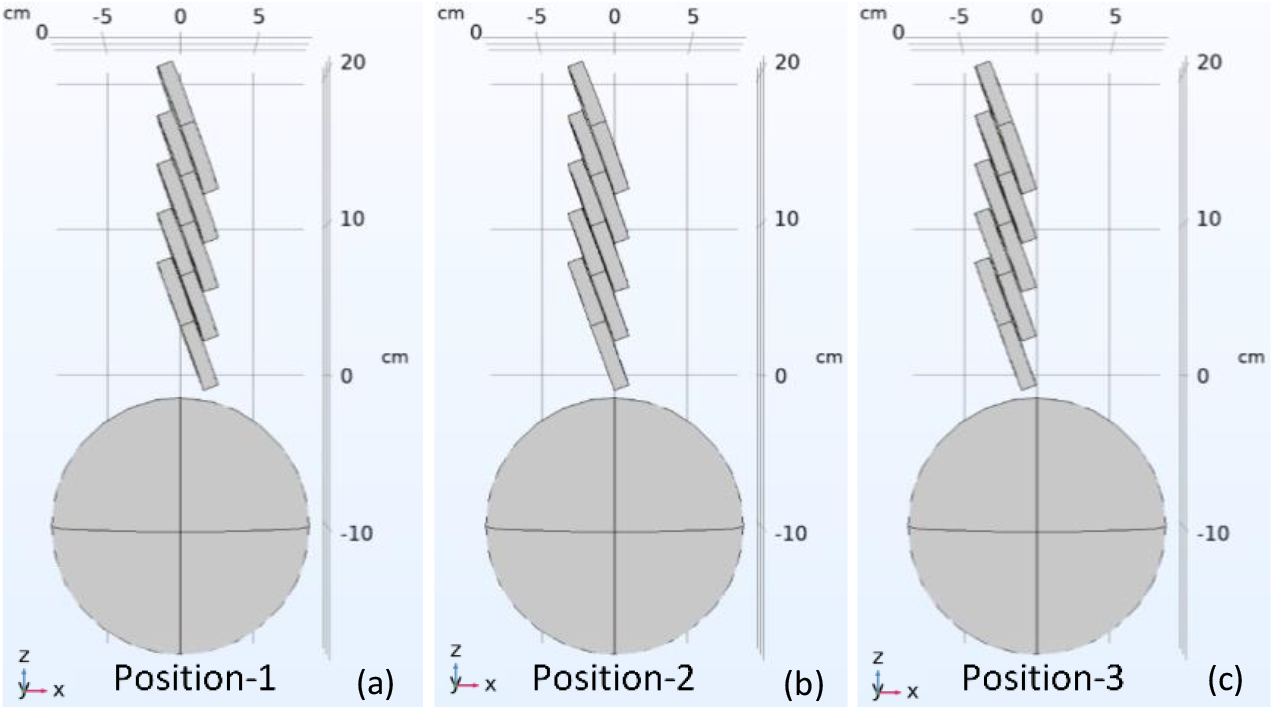
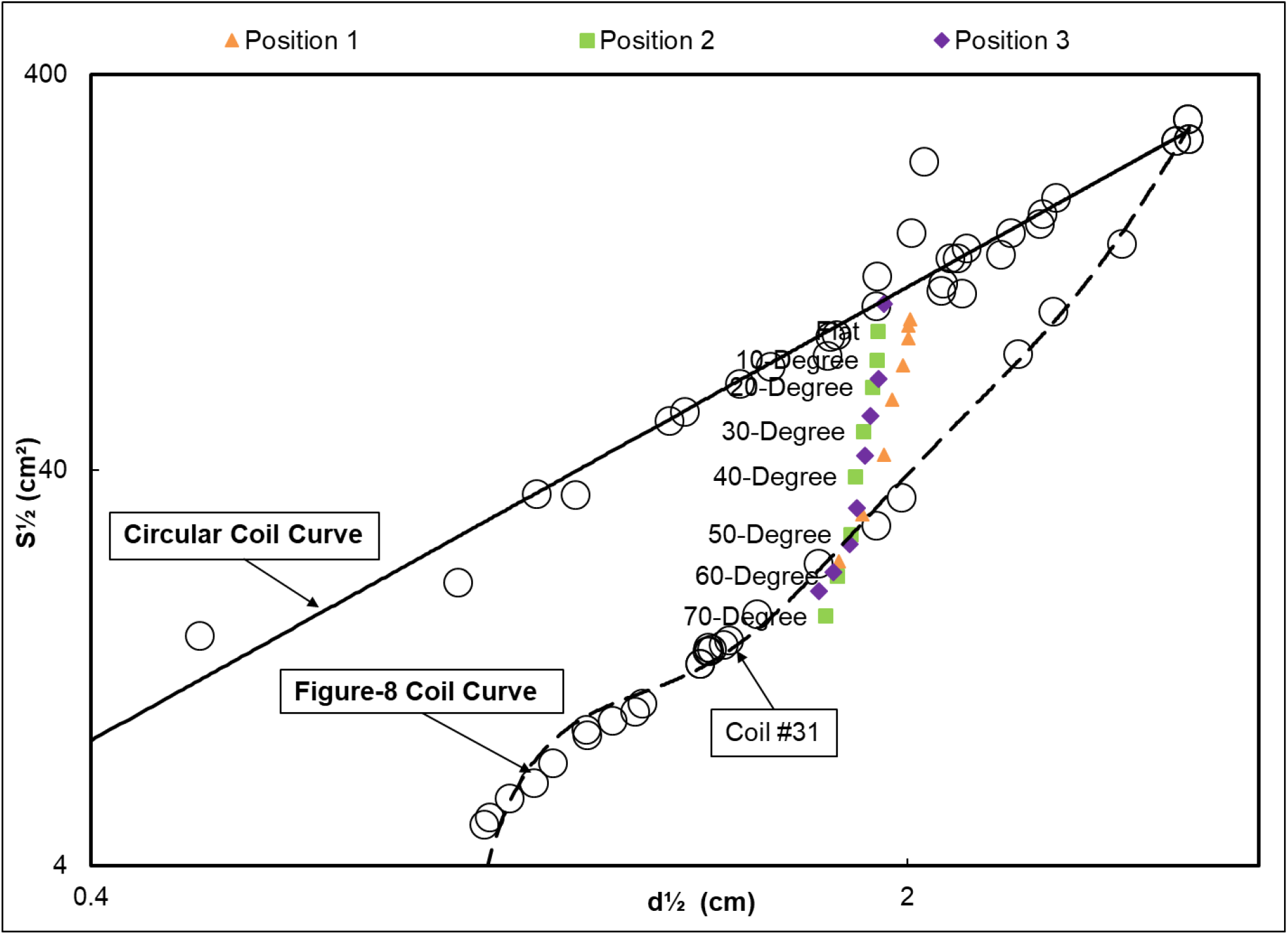
The AT coil position relative to the head model, **a) Position 1:** central axis of the coil is aligned with the center of the head model, **b) Position 2:** the lowest point of the coil is aligned with the head model on the z-axis, **c) Position 3:** the lowest point of the coil is 1 cm away from the central axis of the head model along the z-axis, **d) Effect of the coil position on S**_**1/2**_ **and d**_**1/2**_. The three coil positions relative to the head model were compared. For each of the datasets, the top and bottom data points represent a tilting angle of 0 degrees and 70 degrees, respectively, with 10-degree steps between them. The previously obtained data from [27] is shown in the background for comparison purposes. The inner and outer diameters of the simulated coils are 8 cm and 9 cm, respectively.

Rotating the coil to adjust the coil direction relative to the head model can also improve the depth– spread performance, which is not shown here since the improvement is not significant. From this study, we can understand that the combination of coil stacking and angle tuning has produced an elliptical electric field profile instead of a spherical distribution. If the coil is appropriately aligned with the head model, the induced electric field occupies a smaller half-value volume.

To obtain the tradeoff curve, we vary the coils’ outer diameter between 2 cm and 100 cm while keeping the winding width constant at 1 cm. The tilting angle was fixed at 70 degrees, and the rotation angle was 20 degrees with the lower edge of the coil aligned with the head model’s central axis. Figure 4 shows the results of the depth–spread performance of these coils as a function of coil diameter. The AT coils have a smaller footprint and demonstrate a better tradeoff curve than conventional figure-8 coils. For example, an AT coil with an inner and outer diameter of 3.5 cm and 4.5 cm, respectively, establishes the same d_1/2_ as the figure-8 coil (coil #31) with a 10% smaller spread. When the coil’s inner and outer diameters reach 8.0 cm and 9.0 cm, respectively, the half-value depth experienced a 20% increase compared to the figure-8 coil (coil #31) with the same spread (S_1/2_). This is a considerable improvement compared to the existing figure-8 coils, given that it only occupies about half of the footprint of the corresponding figure-8 coils. The larger coils illustrated in Figure 4 are analyzed for comparison purposes and are not feasible for brain stimulation due to their dimension.

**Figure 4:**
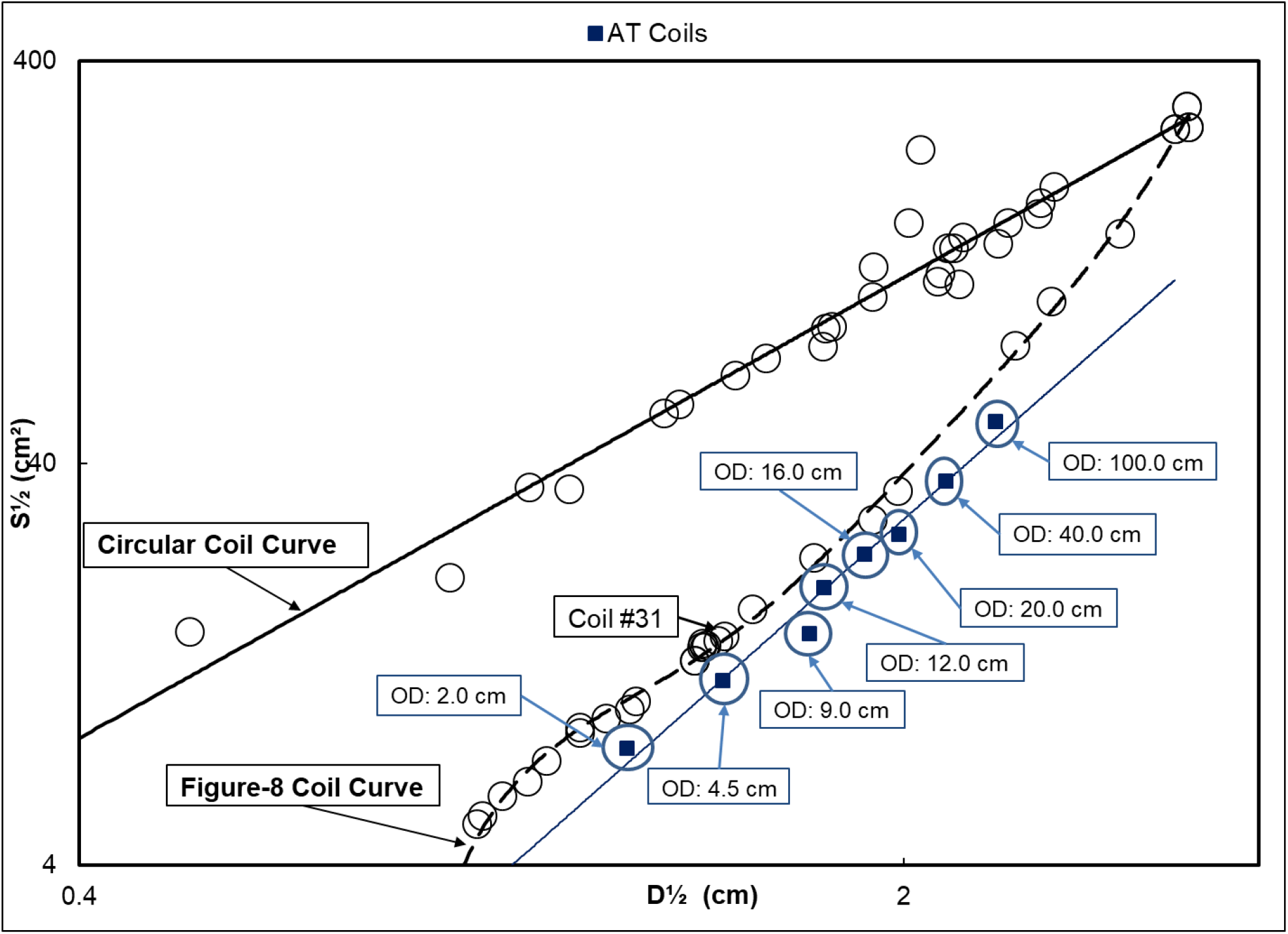
Effect of the coil’s outer diameter on S_1/2_ and d_1/2_. In the plot, the ‘O.D.’ represents the outer diameter of the studied coil. The outer diameter of the AT coils varies from 2 cm to 100 cm, with the winding width kept constant at 1 cm, meaning that the inner diameter is 1 cm smaller than the outer diameter. The previously obtained data from [27] is shown in the background for comparison purposes.

The AT coils have demonstrated a significant potential for multisite brain stimulation. Figure 5 illustrates the proposed apparatus and the implementation of multisite stimulation. In this design, four 4.5-cm AT coils are placed on a mechanical frame (non-metal holder) in a ring structure. Each coil’s relative 3-D location is adjustable and trackable through the translation and rotation stages located on the frame. These stages can be motorized with stepping motors to track all the relative locations. The designed system is convenient to adapt to individual head differences. It has an enhanced depth-spread tradeoff and a small contact area. We used the 4.5-cm AT coils for this design since they establish a smaller spread with a smaller footprint in comparison to conventional coils; the conventional 70-mm figure-8 coil (#31) and the 4.5-cm AT coil have the same half-depth with a 10% smaller spread and 95% smaller footprint for the single AT coil, as shown in Figure 5(b). The smaller footprint and contact surface of the AT coils, in addition to the enhanced depth-spread tradeoff, create flexibility with the coil location and movement relative to the head and the number of coils on the scalp in comparison to conventional dual-coil systems. It should be pointed out that the mechanical frame should not be counted toward the footprint of the apparatus as it is a supporting structure for performing multisite stimulation.

**Figure 5:**
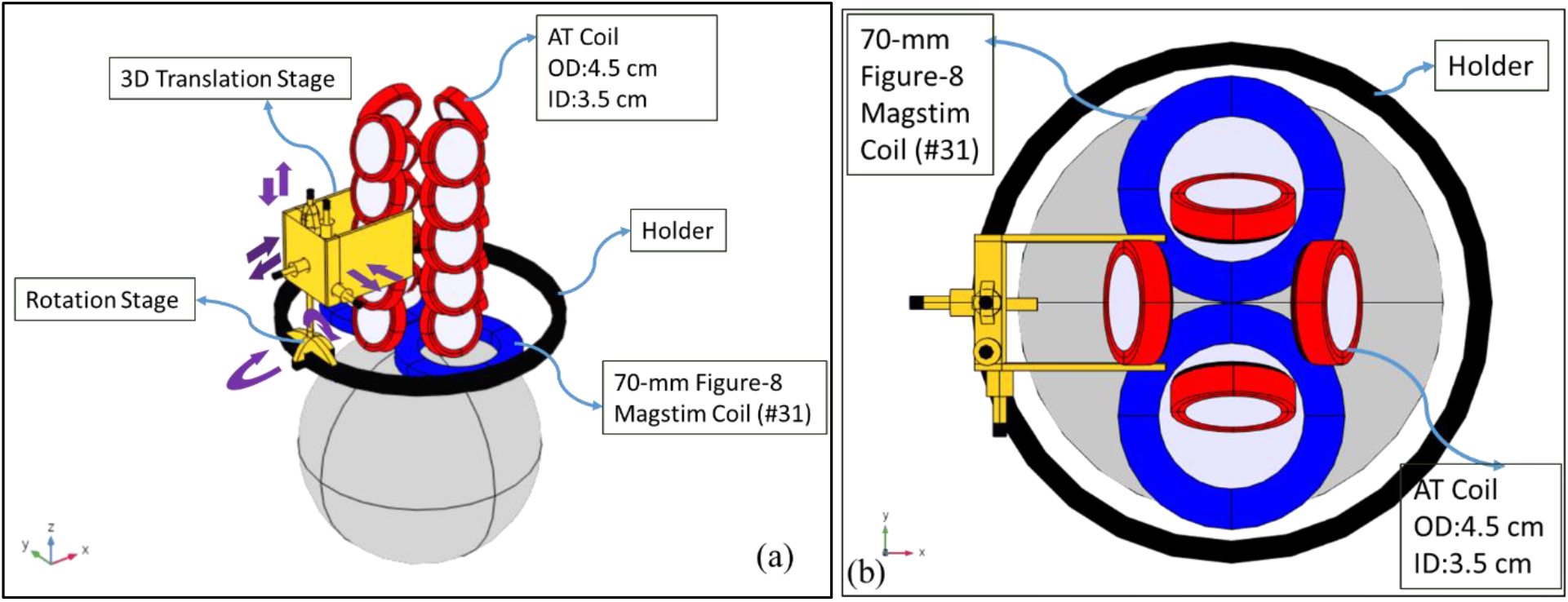
Multisite stimulation apparatus using AT coils, **a) 4 AT coils used for simultaneous stimulation of different brain regions;** A mechanical frame in a ring structure is considered while the location of the coils can be adjusted in this frame, **b) Plan view of the apparatus**. This figure demonstrates the smaller footprint of single AT coils (shown in red) in comparison to the figure-8 coil (shown in blue). For the figure-8 coil, the footprint area is about 120 cm^2^, as shown in Figure 1(c). For a single 4.5-cm OD AT coil, with the same half depth and 10% smaller spread, this value is about 5.5 cm^2^ showing a 95% smaller footprint compared to the figure-8 coil.

Besides using multiple AT coils to build multisite stimulation tools, we can use them as fundamental building blocks for coils with more complex structures and better depth–spread performance^46^. A single AT coil is not symmetric, and it can occupy more V_1/2_ in the head model. For example, adding another AT coil to form a pair and having two AT coil pairs with opposite polarities can create a symmetric structure and produce a more elliptical field distribution. By adjusting various angles among these pairs, we can further optimize the depth–spread performance and obtain an even better depth–spread tradeoff curve. Figure 6 shows an implemented example using this concept. The arrangement of the AT coils in the proposed structure is inserted in the top left corner. In this design, two 80-degree tilted coils with an angle of 20 degrees between them form a pair; the proposed coil design comprises two of these pairs with opposite polarities and an internal angle of 60 degrees between them, and the coil I.D. changes from 3.5 to 29 cm with a 1cm winding width. This result indicates a further improvement of the S_1/2_ and d_1/2_ and confirms AT coils’ role as building blocks for complex coil structures. Compared with the commercial 70-mm figure-8 coil (coil #31), the 3.5 cm inner diameter 4-AT-coil module demonstrates a 10% smaller spread and a 30% larger d_1/2_. Also, the design with a 5 cm inner diameter 4-AT-coil module demonstrates a 15% larger spread and a 50% larger d_1/2_ compared to the corresponding figure-8 coil.

**Figure 6:**
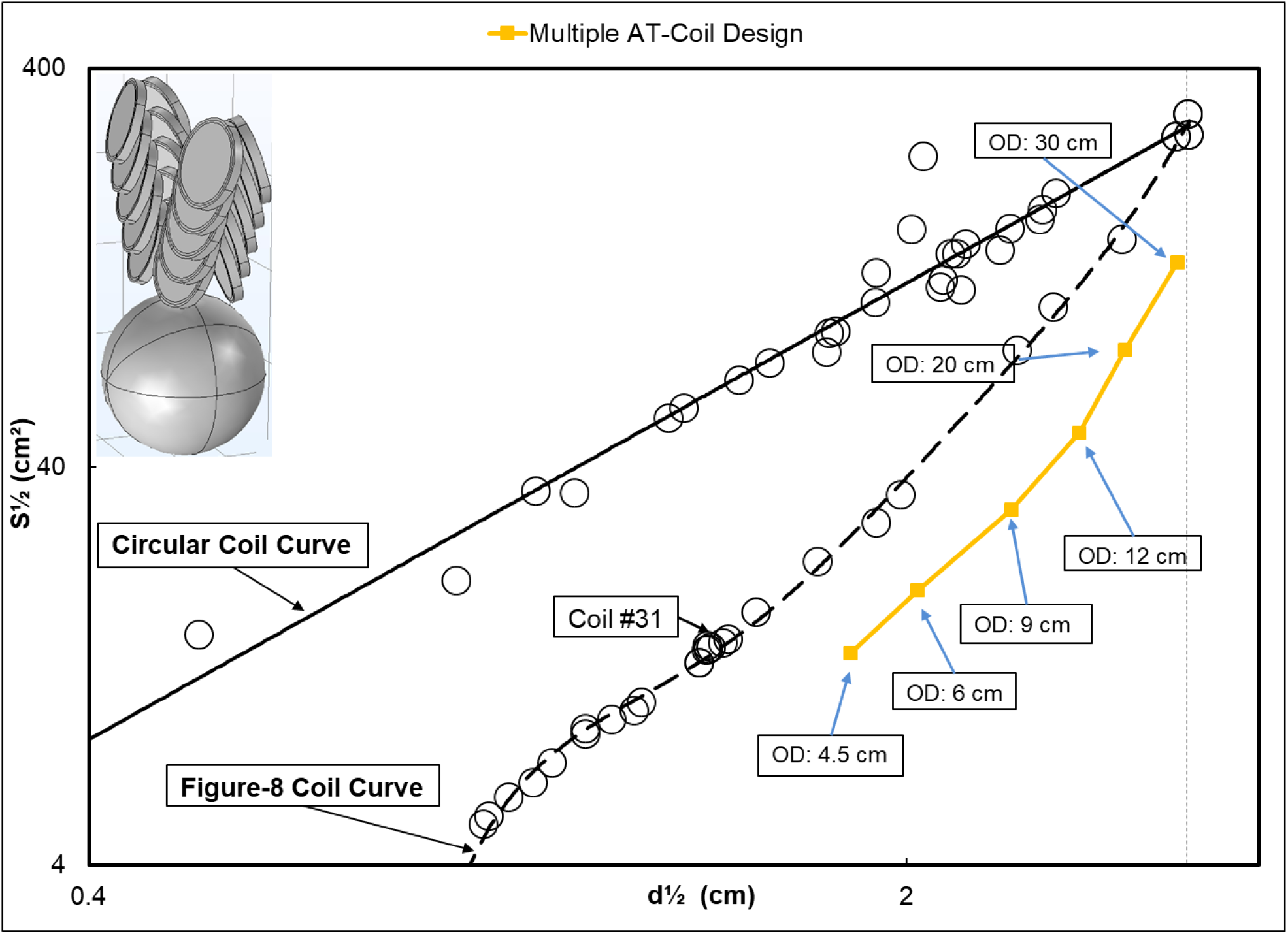
Effectiveness of using Multiple AT coils as pairs for improving the depth–spread. The design of the coil is shown in the top left corner. In the plot, the ‘O.D.’ represents the outer diameter of the studied coil. In this design, two 80-degree tilted coils with an internal angle of 20 degrees form a pair; the proposed coil design is comprised of two of these pairs with opposite polarities and an internal angle of 60 degrees, and the coil O.D. changes from 4.5 to 30 cm with 1 cm winding width. The previously obtained data from [27] is shown in the background for comparison purposes.

### 3.2 EXPERIMENTAL MEASUREMENTS OF INDUCED ELECTRIC FIELD

To verify the simulated data, a total of 8 coil prototypes were fabricated with two different dimensions, two different winding layers, and different tilting angles. The electric field distributions were measured for each of the AT coils and the commercial 70 mm figure-8 Magstim coil using calibrated high-spatial-resolution vector-field probes^50^. The first coil, named the’ Coil- A,’ has an inner and outer diameter of 1 cm and 3 cm, respectively, with nine winding layers. The tilting angle ranges from 0 to 60 degrees with a step of 10 degrees.

The second coil, named the’ Coil-B,’ has an inner diameter of 3 cm and an outer diameter of 9 cm with six winding layers with a tilting angle of 40 degrees. For comparison, both COMSOL simulations and experimental measurements of the electric field decay rate and stimulation hot spot area were conducted.

Figure 7 shows FEM simulation and experimental measurement results of electric field distributions at 1.5 cm away from the surface of these coils in air. The results indicate that for all the proposed AT coils, the maximum electric field’s location is relatively the same, close to the coil’s tilted edge, while for figure-8 coils, it is happening in the center of the coil. This observation further proves the usefulness of the AT coils for multisite stimulation.

**Figure 7:**
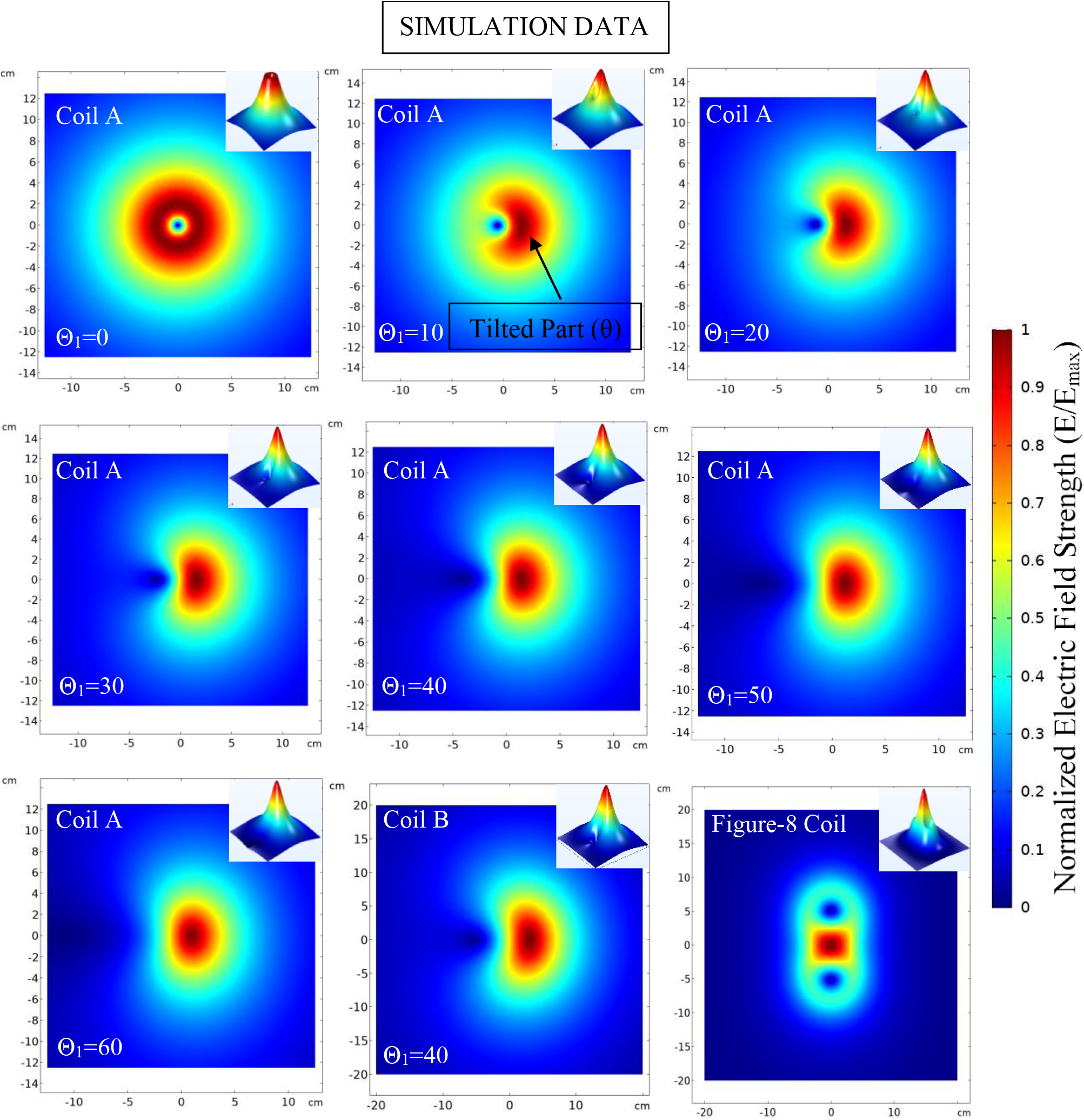
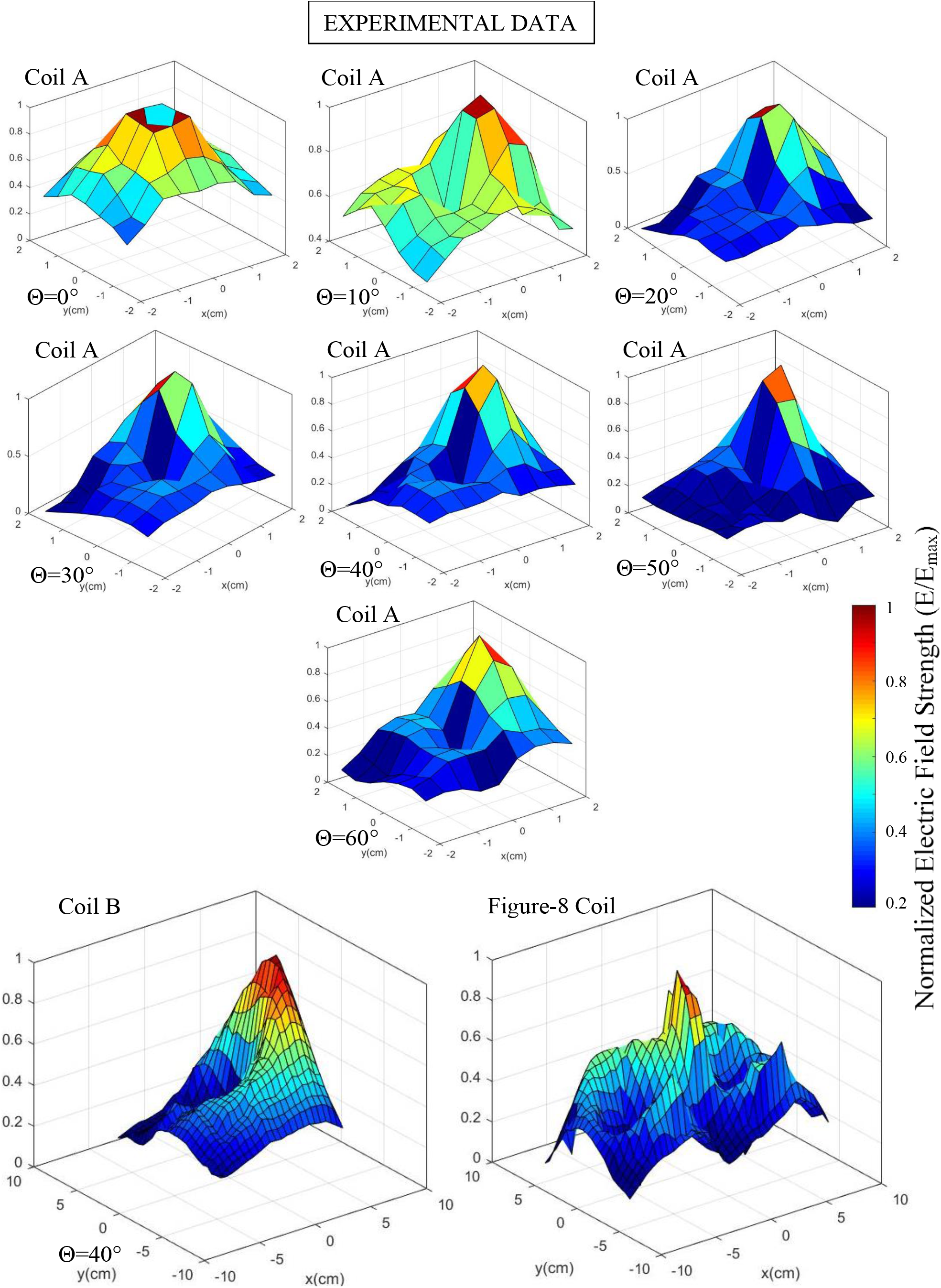
Electric field distribution for the proposed coils with different tilting angles based on the simulations and experimental measurements; The simulations demonstrated the 2-D and 3-D distribution of the electric field. All the experimental measurements were performed in 1.5 cm distance from the coil’s surface with 5-mm lateral steps using calibrated high-spatial-resolution vector-field probes. The dark red areas show the highest intensities encountered for both simulations and experiments; all the data is normalized to the highest electric field intensity. In the experimental data, the scanned area for the coil-B and the figure-8 coil is larger than the coil-A series due to their larger size, which resulted in more pixels for the two coils.

Although it is difficult to measure the S_1/2_ directly, one way to represent the spread properly is to check the size of the hot spot as defined in the following procedure. We scan and obtain the maximum electric field strength at a fixed distance away from the coils. At each distance, hot spot size is defined by measuring the area with the electric field strength above a selected percentage of that measured maximum strength. For example, at the distance of 1.5 cm away from the coils, we define our hot spot size as the areas with an electric field intensity of more than 90% of the measured maximum electric field strength. Although we can choose to set lower percentages, we specified a greater than 90% to avoid scanning a larger area for a smaller percentage without losing fairness and accuracy.

As shown in Figure 8 (a), tilting the coil’s wrapping angle significantly decreases the focal spot size. At a 10-degree tilted angle, the spot size drops 80% from that of the flat coil. The spot size further reduced an additional 70% from around 5.0 cm^2^ for the 10-degree coil to about 1.5 cm^2^ for the 60-degree coil. The focal spot size of the entire coil-A series is smaller than that of the figure-8 coil. On the other hand, the 40-degree tilted coil-B has a slightly bigger focal spot area than the studied figure-8 coil, which can be overcome if higher tilting angles are used. The argument can be proved by using the coil-A series as an example. The hot spot size reduced 15% from a 40-degree coil to a 60-degree coil. If the same effect is applied, the B coil’s hot spot size becomes smaller than that of the figure-8 coil.

**Figure 8.**
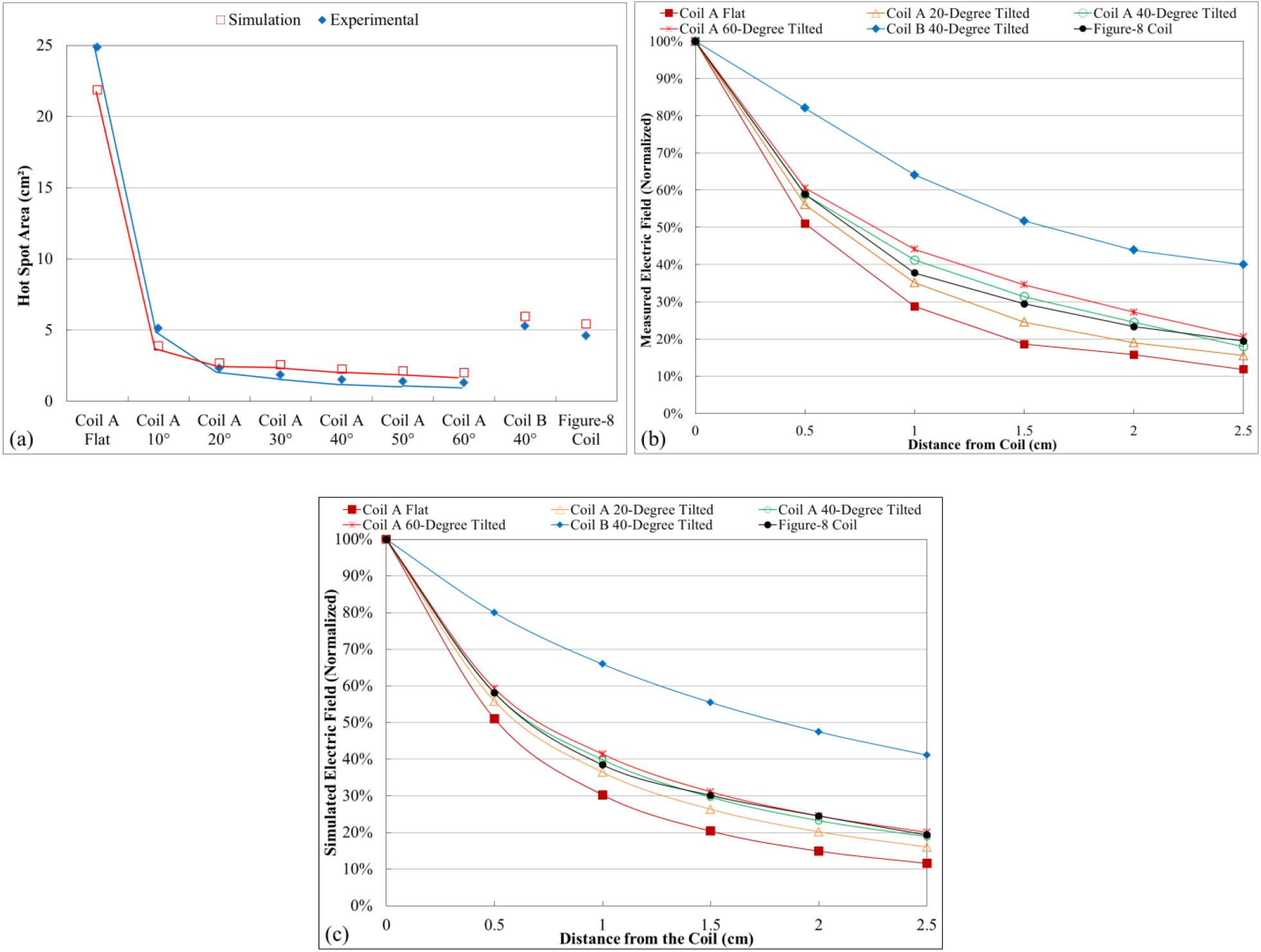
**a) Hot spot size for experimental data and simulations**. The hot spot size is measured and simulated at 1.5 cm away from the surface of the coil, **b) Electric field intensity decay rate (in percentage) as a function of depth for the experimental measurements, c) Electric field intensity decay rate (in percentage) as a function of depth for FEM simulations**. The presented distance in this Figure is measured from the surface of the coil in the air. In contrast, in Deng et al.^27^, the distance is considered from the cortex, which is 2 cm away from the coil’s surface. Considering the same concept for this figure (from 2 cm to 2.5 cm) indicates about 10% improvement from the figure-8 coil to Coil B, which matches the data obtained in the first part of this study in Figures 2 through 4.

The electric field intensity decay rates based on the experimental data and simulations are shown in Figures 8 (b) and 8 (c), respectively. We first normalized the measured data for an accurate comparison. As shown in these plots, the decay rates gain improvements by increasing the tilting angle. At a depth of 2.5 cm from the coil, the flat coil’s decayed remaining value is less than 15.0% for both experimental and simulation data. In contrast, for the 60-degree tilted coil, the value increases to 25.0%, indicating an improvement of the decay rate. It can also be observed that the 40-degree tilted coil B has a much slower decay rate than the figure-8 and all other coils. The depth-spread performance of the AT coils is summarized in Table 1. The hot spot area of A coils is smaller than the figure-8 coil, while the B coil’s hot spot area is comparable to the figure-8 coil. Nevertheless, its decay rate is more than twice slower than the figure-8 coil. More than 40% of field strength remained at 2.5 cm distance for the B coil compared with the figure-8 coil with less than 20% left. Additionally, the simulation data in Figure 2 shows the improvement in performance for higher tilting angles compared to the 40-degree tilted coil.

**Table 1.** The hot spot area and the induced electric field intensity decay rates from experimental measurements and FEM simulations.

| Coil Type | Tilting Angle (°) | Hot Spot Area (cm<sup>2</sup>) |  | Remaining % at 2.5 cm |  |
| --- | --- | --- | --- | --- | --- |
|  |  | Measured value | Simulated value | Measured value | Simulated value |
| Coil A | Flat | 24.88 | 21.89 | 11.8 | 11.6 |
|  | 10 | 5.13 | 3.9 | 13.6 | 14.1 |
|  | 20 | 2.34 | 2.7 | 15.6 | 16.1 |
|  | 30 | 1.85 | 2.57 | 16.7 | 17.9 |
|  | 40 | 1.52 | 2.28 | 17.9 | 18.9 |
|  | 50 | 1.41 | 2.13 | 19.0 | 19.6 |
|  | 60 | 1.3 | 2.01 | 20.5 | 20.1 |
| Coil B | 40 | 5.28 | 5.98 | 40 | 41.1 |
| Figure-8 |  | 4.61 | 5.44 | 19.4 | 19.3 |

One particularly notable fact is that for the 50- and 60-degree tilted angles, the small diameter A coils can accomplish better decay rates than the much larger diameter figure-8 coil. This performance shows that the proposed design can produce elevated electric field intensity in deeper brain regions due to the field redistribution, or more precisely, focusing effect through angle tuning.

## 4. DISCUSSION

The proposed coil design has the main advantage of occupying a much smaller contact surface during the stimulation due to its vertical stacking. For multisite brain stimulation, only one power supply circuit unit is required for the AT coils. Since all the elements are the same with an equal inductance, the combined inductance can easily be adjusted with parallel and serial connections to further improve depth-spread performance and multisite stimulation. The approach of using a uniform building block to construct a composite coil structure provides other benefits. First, mass production of identical small units can help to reduce cost and increase quality. Second, it provides flexibility in designing and implementing a novel generation of TMS tools by merely adjusting the relative geometric locations of these identical building block coils. The single AT coils used in the composite structure can also be adjusted with the tilting angle and the stacking number to match the required stimulation results. Third, the AT coils’ simple design allows for easier replacement of possible defective elements in the multi-coil apparatus versus repairing or replacing the whole unit like other reported complicated multisite stimulation structures. Noticeably, compared with all the reported or existing complex coil structures designed for deep brain stimulation, our simple 4-uniform-AT-coil design has better depth-spread performance, as shown in Figure 6. This novel coil design provides a promising new generation of future high depth-spread performance and multisite TMS tools.

The proposed coils’ energy consumption was analyzed based on the neural stimulation threshold of 100 V/m^51^ at 3 cm from the head’s surface using the magnetic flux density and magnetic field intensity vectors^52^. The analysis indicated that the required energy to reach the threshold increases exponentially by reducing the coil’s diameter size. For example, compared with the 70-mm figure-8 coil (coil #31), the 9cm-OD AT coil demonstrates the same spread, a 20% larger penetration depth, and an 80% less footprint while the energy required to reach the threshold for the figure-8 coil is 70% less. This result is expected since the 9cm-OD AT coil has only a 20% footprint of that of the figure-8 coil (coil #31). Considering a figure-8 coil with the same footprint as the 9cm-OD AT coil, it exhibits about 25% higher energy consumption but with much worse depth performance.

## 5. CONCLUSION

We have proposed a stacked and tilted ring structure TMS coil, which has a less than 50% smaller footprint and better depth–spread performance than a conventional figure-8 coil. Based on a similar principle, in our earlier works, we have introduced a rodent coil that can provide an activation spot size of less than 2 mm and demonstrated single limb motor cortex activation in rodents. In this work, we offer a comprehensive picture of the depth–spread improvement by using angle-tuning to control the spread and use coil stacking to increase depth penetration. We have shown, theoretically and experimentally, that the AT coils have improved field decay rates to reach deeper regions with a higher field intensity and a reduced field spot size compared with conventional figure-8 coils of comparable spread. The proposed coil can serve as a fundamental building block for more complex coil structures to further improve depth–spread performance.

## ACKNOWLEDGEMENT OF FUNDS

The research was supported by the NSF grant ECCS-1631820, NIH grants MH112180, MH108148, MH103222, and a Brain and Behavior Research Foundation grant. It was also partly supported by the Intramural Research Program of the National Institute on Drug Abuse, National Institutes of Health.

